# The evolution of function in the DNA binding domain of the CRP/FNR family

**DOI:** 10.1101/2025.07.04.663149

**Authors:** Madhumitha Krishnaswamy, Aswin Sai Narain Seshasayee

## Abstract

Transcriptional regulation, facilitated by transcription factors (TFs), contributes to bacterial response to environmental and cellular perturbations. How have TFs diversified and gained their functions? We use the CRP/FNR family to study this question across ∼6000 prokaryotic genomes. Characterized by homology to TFs CRP (cyclic AMP Receptor Protein) and FNR (Regulator of Fumarate and Nitrate Reductase) in *Escherichia coli*, the two functions of this family – sequence-specific DNA binding and transcriptional activation by direct contacts to RNA polymerase - are conferred by the helix-turn-helix containing DNA binding domain. We constructed a rooted phylogeny of this domain and performed residue conservation analysis on extant and in-silico reconstructed ancestral sequences of CRP/FNR family members. Residues that confer DNA binding specificity likely emerged in the overall ancestor of the sequence family. This ancestor does not fall into classes containing either *E. coli* CRP (Eco-CRP) or *E. coli* FNR (Eco-FNR) clades, but instead a large and diverse class that we call “CRP-like”. Residues key for base-specific DNA binding remain largely conserved in the family. Residues (together called AR1) which make direct contacts with αCTD of RNA Polymerase are conserved only in the Eco-CRP class and restricted to *Gammaproteobacteria*. The corresponding ‘287 determinant’ interface on αCTD is again fully conserved only in *Gammaproteobacteria*, which contain Eco-CRP members, following step-wise evolution initiated at the Proteobacterial common ancestor. These suggest suggesting co-emergence of the interface residues in Eco-CRP. Our analysis hence shows that Class I transcriptional activation through AR1 on CRP and the determinant of αCTD, as described in *E. coli* CRP, is phylogenetically restricted, while base-specific DNA binding has been present in the CRP/FNR family throughout its evolutionary history.

## Introduction

Transcription factors (TFs) regulate gene expression, typically by regulating transcription initiation (Browning & Busby, 2004). However, studies of minimal cellular gene content indicate that TFs are not essential for the basic functioning of an organism (Gil et al., 2004). When did TFs emerge and how have they evolved? The evolution of TF function, presumably from a non-specific DNA binding ancestor (Visweswariah & Busby, 2015), would require a DNA binding protein to gain specificity for recognizing certain DNA sequence motifs; for TFs that activate transcription, an interface for recruiting RNA polymerase; and for TFs that respond to a cellular signal, the ability to allosterically transduce the presence of the signal to DNA binding activity. In a recent work, we provided evidence to suggest that sequence-specific DNA recognition in the TF IHF evolved from an ancestor resembling its non-specific DNA binding relative HU. This evolution appears to have been achieved by repurposing pre-existing residues as well as by evolving new residues (Nandy et al., 2025).

The CRP/FNR family is a well-characterized TF family that contains 2 global regulators, among other proteins: CRP (cyclic AMP Receptor Protein) and FNR (Regulator of Fumarate and Nitrate Reductase). This family of TFs is characterized by a DNA-binding helix-turn-helix domain and a second domain that senses a signal (cyclic AMP in CRP and redox states in FNR). The helix-turn-helix domain in the *E. coli* CRP enables two abilities: (a) sequence-specific DNA binding and (b) an interface for binding to the α C-Terminal Domain (αCTD) of the RNAP, which allows the TF to activate transcription by what is termed Class I activation (Busby & Ebright, 1999; Lee et al., 2012). Sets of residues involved in these functions have been studied in great detail in *E. coli* CRP (Lawson et al., 2004). Further, *E. coli* CRP binds to the genome at hundreds to even thousands of sites, regulating proximal gene expression at some but not at others (Grainger et al., 2005). Thus, it has been proposed that CRP might function not only as a TF but also as a chromosome-shaping protein. Could this protein be an evolutionary intermediate between a classical chromosome-shaping non-specific DNA binding protein and a sequence-specific TF (Visweswariah & Busby, 2015)? These biochemical and logistical advantages of CRP, and the availability of the genomes of thousands of bacterial species allows us to investigate, within a phylogenetic framework, the evolutionary traectory of both the DNA binding and RNA polymerase-recruiting functions, which are well defined in the helix-turn-helix domain of *E. coli* CRP, in the broader family of the CRP/FNR family of TFs.

In this study, we use protein sequence and phylogenetic analyses to ask how evolution has built the helix-turn-helix domain of *E. coli* CRP to specifically bind DNA and contribute to class I transcription activation by recruiting RNA polymerase through an interface with its αCTD.

## Materials and methods

The dataset of 6,116 prokaryotic genomes used follows from (Nandy et al., 2025), along with the corresponding genus level curated 16S rRNA species tree (N=1454).

### Protein curation and ortholog classification

hmmsearch (Eddy, 2011) was used to identify all proteins containing both the ligand binding domain (Pfam: PF00027) and the helix-turn-helix-containing DNA binding domain (Pfam: PF13545) as defined for the CRP/FNR family of proteins, resulting in a final dataset of 13,603 domain sequences (see Supplementary Methods). A classification based on multiple clustering (based on the bitscore values between protein ortholog pairs) methods and functional annotations (Kegg Orthology IDs of resulting hits) resulted in the identification of 10 different classes of the helix-turn-helix DNA binding domains in this family (see Supplementary Methods).

### Principal Component Analysis and NJ tree

Domain sequences were clustered using (a) Principal Component Analysis or PCA (Konishi et al., 2019) (See Supplementary Methods) and (b) Neighbour-Joining Tree calculations done using FastTree (Price et al., 2010) with the -noml option, which skips the NNI and ML calculation steps in tree construction.

## Gene tree

### Domain alignment and curation

The protein sequences were first trimmed to their domain coordinates as obtained from Pfam. The trimmed domains were then aligned using mafft (Katoh & Standley, 2013) -auto with a gap extension penalty of -0.123, identified as being an optimal penalty value to improve alignments (Long et al., 2016). As these domain sequences will be used to construct a codon alignment-based gene tree, traditional trimming methods of sequences could not be easily employed; instead, a pruning/exclusion method was employed. First, extremely long sequences are removed from our dataset. Following this, sequences contributing to a large portion of gappy regions in the domain are removed followed by multiple iterations of re-alignment to arrive at a reasonable alignment. (See Supplementary Methods). This alignment was used for conservation statistics prior to tree-based conservation analysis.

### Tree parameters

To construct a gene tree, the protein alignment was first converted to a codon alignment using PAL2NAL (Suyama et al., 2006), by giving the protein alignments along with their corresponding CDS gene sequences as input. ModelFinder (Kalyaanamoorthy et al., 2017) was then run with a sampled alignment using IQ-TREE 2 (Minh et al., 2020) to identify the best model (KOSI07+F+R4) to be used for the construction of the complete gene trees. Trees were constructed with 1000 ultrafast bootstraps (Hoang et al., 2018). Domains belonging to the same homologous superfamily are evolutionarily related and structurally conserved and could serve as good outgroups. Under this assumption, all sequences belonging to the “Winged-helix-turn-helix superfamily” were examined against alignments of sequences sampled from our dataset until a successful outgroup (a fungal sequence containing Pfam: PF01035), which gave consistent results upon repeated subsampling during tree construction, was identified. All the constructed trees were annotated and visualized using iTOL (Letunic & Bork, 2024).

### Ancestral sequence reconstruction

A gene tree of the helix-turn-helix domain was constructed using IQ-TREE 2 with a specified outgroup (−out). This tree was used to infer ancestral sequences (−asr) by maximum-likelihood. The “state” file output from IQ-TREE 2 was processed using an in-house script to give the ancestral protein sequences for each node. Key nodes identified from the gene tree were appended to the domain alignment and used for further analysis.

### Phylogenetic Reconciliation

Phylogenetic reconciliation of a species tree and gene tree is usually performed to identify the nodes (on the species tree) which served as sites for gene duplication, loss or transfer by comparing the tree topologies of a species and gene tree and for multiple iterations, arriving at the most optimal of events. In this case, the 16S rRNA tree, constructed in previous work (Nandy et al., 2025), and the appropriately pruned gene tree were reconciled using Ranger-DTL (Bansal et al., 2018) to estimate the number of duplications, transfers and speciation events. 100 reconciliations performed with default parameters were combined using Aggregate Ranger to give a final mapping of nodes experiencing duplication, transfer and speciation events. OptRoot by Ranger-DTL uses phylogenetic reconciliation to arrive at a rooting of the gene tree that minimizes the overall costs of events and hence, is used to identify extant sequences that could serve as possible roots. OptRoot was run 100 times over different threshold transfer costs (distance dependent), and the results were summarized using SummarizeOptRootings.

#### Residue conservation analysis

For initial conservation analysis, Jalview (Waterhouse et al., 2009) provided a set of properties which was then used to calculate conservation statistics based on the multiple sequence alignment.

The PSSM (Position specific scoring matrix) scores for a reference set of residues used to evaluate conservation are computed using the same methods as in our previous work (Nandy et al., 2025). The reference sequences and sites used for the calculation of the log odds matrix that forms the PSSM were derived from the literature for binding with αCTD and DNA, as well as key determinants identified for αCTD. These papers are cited where appropriate in Results. Residue positions on the multiple sequence alignment were identified based on the reference, and the sequences in the alignment were scored using the corresponding sites in the alignment.

In addition, a BLOSUM based scoring system was used to compute conservation of key residues. In this method, the standard BLOSUM62 matrix for amino acid substitutions was used instead of the log odds matrix computed from the reference sequence as follows:

The BLOSUM62 substitution matrix, accounting for a gap penalty of -0.12 is represented as a 21*21 matrix. For a reference set of key residues of length *n*, the BLOSUM62 (termed *B)* matrix to be used becomes a *n*\*21 matrix. For each residue “*i*” in the set *n*, the *i*^th^ row of *B* represents all the substitution scores based on the 21 possibilities at each position (20 residues + gap). Matrix *B* is used to score each sequence in the alignment at the corresponding residue positions, giving a BLOSUM62 score for the set of key residues.

Since similar results were obtained using both the log-odds matrix and the BLOSUM scores, all further references to PSSM refers to the scores computed from the log-odds matrix.

## Results

### Ancestry of the DNA binding domain of the CRP/FNR family

The TF CRP from *Escherichia coli* contains a DNA-binding helix-turn-helix domain and a ligand-binding domain that binds cyclic AMP (cAMP). This domain architecture is also shared by another global TF, FNR, in which the ‘ligand-binding’ domain does not bind cAMP but instead contains an iron-sulphur cluster making it a redox sensing transcription factor (Spiro, 1994). Here we refer to the helix-turn-helix family present in CRP and FNR as “c-f-HTH” for convenience; note however that the helix-turn-helix itself is a much larger superfamily found across a wide variety of DNA binding proteins. The c-f-HTH domain is responsible for specific DNA recognition and also contains the so-called Activating Region 1 site (AR1). The AR1 forms an interface with the α-C-terminal domain (αCTD) of the RNA polymerase thus recruiting it for transcription (De Crombrugghe et al., 1984). To understand how specific DNA binding and RNA polymerase recruitment have emerged and evolved within the helix-turn-helix domain in this classical transcription activator family, we analysed ∼13,500 c-f-HTH domain sequences from across ∼6,000 prokaryotic genomes. We found that while proteins containing the c-f-HTH domain are present in both Archaea and Bacteria, those with both the c-f-HTH and the ligand binding domain are present only in Bacteria, with a notable absence in the clade containing Candidate Phyla Radiation (CPR group).

Using various sequence classification approaches and databases, we classified the proteins in our dataset into 10 groups as described in Supplementary Methods (Supplementary Figure 1). Briefly, we use the term “Eco-CRP” (N=376) to refer to a particular clade of proteins that includes *E. coli* CRP; and “Eco-FNR” (N=661) for the corresponding clade containing *E. coli* FNR. We also identified a large group of sequences annotated as CRP in orthology databases but are distinct from Eco-CRP; these we call as “CRP-like” (N=2,841). We similarly defined a group of “FNR-like” proteins (N=1,197). Eco-CRP proteins are present exclusively in *Gammaproteobacteria*, while Eco-FNR proteins are present in both *Gammaproteobacteria* and *Betaproteobacteria*, consistent with previous studies (Körner et al., 2003) (Figure 1A).

**Figure 1:**
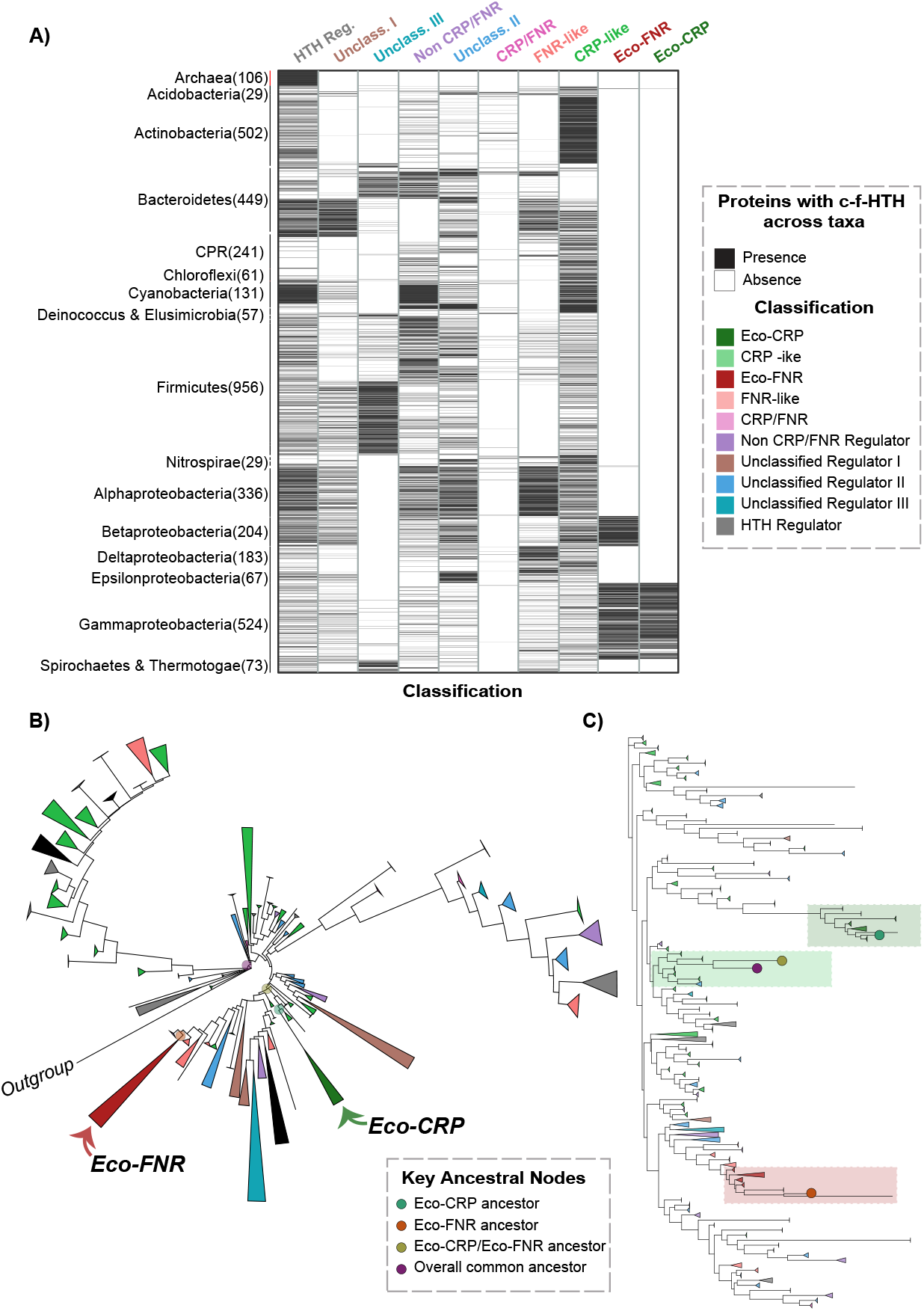
Classification and phylogeny of the c-f-HTH domain. (A) Heatmap showing the presence/absence (black/white) of the different classes across all taxa which showed presence of the c-f-HTH domain (N=4,114), represented at the phylum level (class level for certain diverse phyla). Note - grayscale heatmap with thin binary values. (B) Collapsed gene tree (N=10,224) of the c-f-HTH domain sequences rooted using the DNA binding domain of a methyltransferase enzyme from Fungi. Size of collapsed nodes indicates number of extant tips, and coloured based on classification. Collapsed and visualized using iTOL. (C) Collapsed Neighbour Joining (NJ) tree (N=10,227) of c-f-HTH domain sequences along with 4 key ancestral nodes, as indicated in legend. Clades containing nodes of interest are shaded on the basis of overall regulators present in the clade. Collapsed and visualized using iTOL.

To characterize the ancestral sequences and the evolutionary trajectory of the c-f-HTH domain, we constructed a rooted codon-based gene tree from an alignment of 10,223 c-f-HTH domain sequences (See Supplementary methods). We tested the rooting of the gene tree using multiple methods. By aligning a sample of c-f-HTH sequences with other members of the same protein superfamily, we identified a DNA binding domain of a fungal (*Sporothrix schenckii*) 6-O-methylguanine-DNA methyltransferase enzyme as an outgroup (Figure 1B). As controls, we also used phylogenetic reconciliation to identify other possible rootings (see Methods). The overall topology of the two trees and subsequent major conclusions regarding conservation (Supplementary Figure 2) are consistent.

Using these trees we performed maximum-likelihood-based ancestral sequence reconstruction of the c-f-HTH family sequences at key nodes. The last common ancestor of the c-f-HTH family clusters with neither CRP nor FNR from *E. coli* but instead lies somewhere with the large and diverse set of sequences belonging to the ‘CRP-like’ class. The exact clade of CRP-like to which this ancestor clusters with differs between outgroup-based and reconciliation-based trees (inset, Supplementary Figure 2). However, this difference does not affect the results on the evolution of key functional residues described below. The last common ancestor of Eco-CRP and Eco-FNR together clusters with ‘CRP-like’ as shown in a Neighbour joining tree (Figure 1C), as well as by a PCA (Supplementary Figure 3). As controls, the common ancestor of Eco-CRP lies within the same clade, and that of Eco-FNR clusters with its descendants. Based on the taxonomic distribution of the different classes of proteins we can say that the Eco-CRP/Eco-FNR common ancestor probably emerged at the common ancestor of beta- and gamm-proteobacteria.

### Evolution of key functions of the c-f-HTH domain

To understand the evolution of key functions performed by the domain as described in Eco-CRP, we first define the key residues, taken from the literature including a comprehensive structural analysis of CRP-αCTD-DNA complex in *Escherichia coli* (Benoff et al., 2002; Lawson et al., 2004; Parkinson et al., 1996), which we broadly classify into 3 categories, described as follows (Table 1). The AR1 site (10 residues in *E. coli* CRP) interacts with the ‘287 determinant’ of the αCTD of RNA Polymerase to facilitate transcriptional activation by a Class I mechanism. The residues in the c-f-HTH domain which contact the DNA consensus motif do so in 2 ways: (a) by contact with the phosphate backbone (7 residues in *E. coli* CRP) or (b) by contact with specific DNA base edges (3 residues in *E. coli* CRP).

**Table 1:** Key functional residues for the c-f-HTH domain of CRP/FNR family. Each residue is annotated based on its nature and position as defined for Escherichia coli CRP in (Benoff et al., 2002; Parkinson et al., 1996)

| Functional region | Residues of interest in *E. coli* CRP |
| --- | --- |
| Activating <u>region I</u><br>(AR1) | Pro154, Asp155, Ala156, Met157, Thr158, His159, Pro160, Asp161, Gly162, Met163, Gln164 |
| Phosphate backbone<br>contacts | Lys166, Arg169, Gln170, Ser179, Thr182, Lys188, His199 |
| Specific base contacts | Arg180, Glu181, Arg185 |

AR1 is highly conserved only in Eco-CRP. A distinct set of residues is conserved in Eco-FNR. However, these sites show no conservation whatsoever in other clades of this family including CRP-like and FNR-like. This holds true even when conservation is defined not by the identity of the amino acid residue but by its physicochemical class. In contrast to AR1, residues key for DNA binding show a much higher conservation across all the classes (Figure 2). Residues making DNA backbone contacts show patchy conservation across the classes, with different residues of similar physicochemical types occupying the conserved site at some positions. Residues responsible for base-specific interactions are highly conserved across the phylogenetic tree. Thus, key functionalities of the c-f-HTH domain show different levels of conservation across different orthologous groups within the CRP/FNR family. Specifically, the ability of the c-f-HTH domain to facilitate Class I transcriptional activation through the interface defined in *E. coli* may be restricted to Eco-CRP, unlike DNA binding, which is more widespread.

**Figure 2:**
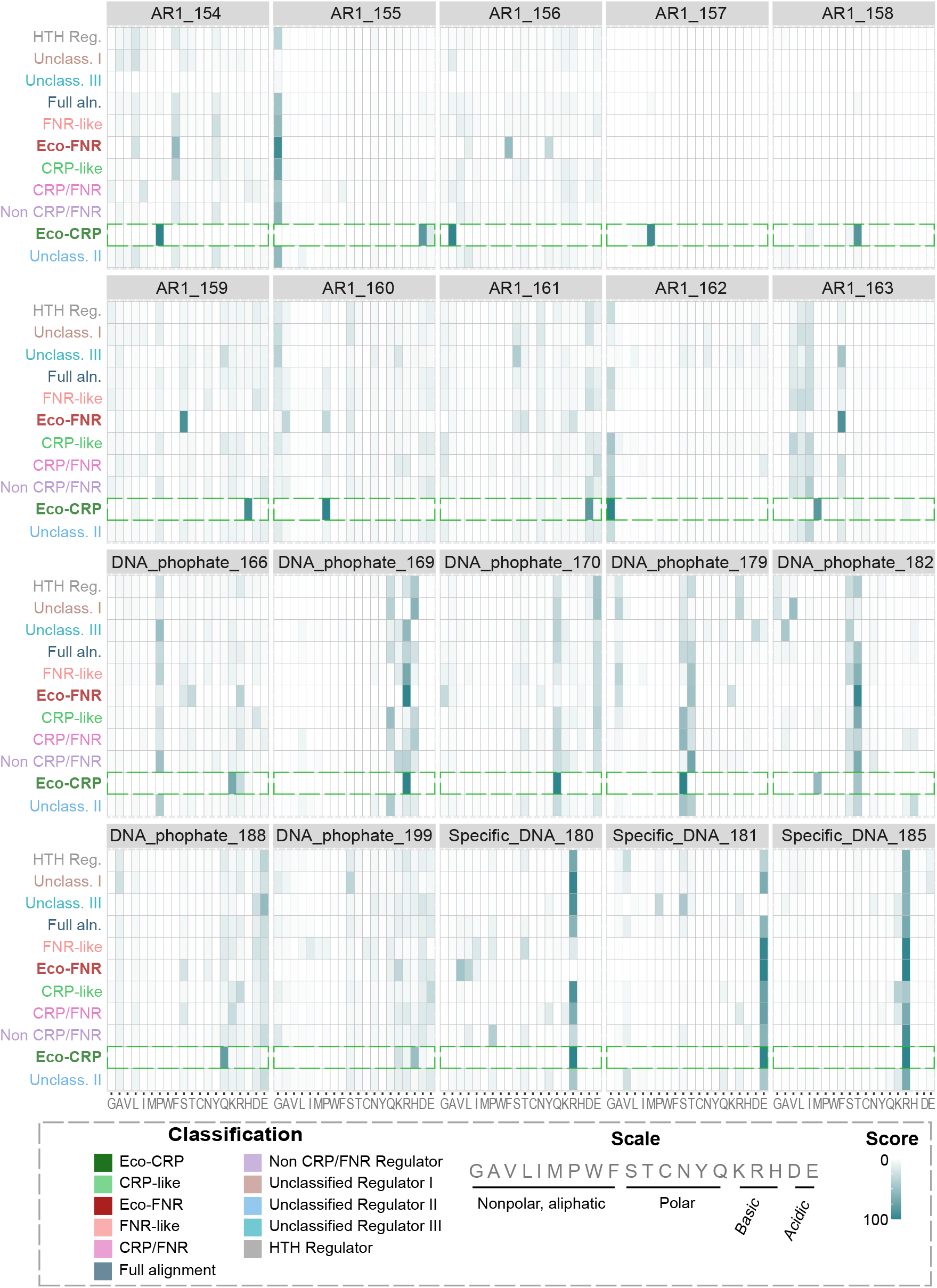
Conservation of key functional residues. Heatmaps for each key residue of interest across the different domain classes, ordered by physicochemical properties. Colour of intersecting boxes is scaled according to the amino acid frequency from low (light) to high (dark) and indicates occurrence of a particular amino acid for the specified key residue in all members of a class. The distributions for Eco-CRP are indicated in the green dashed boxes.

To trace the possible evolutionary trajectory of these functionalities, we analyzed ancestral sequences inferred using maximum-likelihood methods from the gene tree. We computed the PSSM (Position-specific scoring matrix) scores for each of the different categories of key residues in all the ancestral sequences (see Methods). This measures their conservation with reference to the set of residues defined in the literature as necessary for a certain function in *E. coli* CRP. We found that residues of AR1 emerge only in the common ancestor of the Eco-CRP clade and are retained downstream (Figure 3A). Residues which bind to DNA by making phosphate backbone contacts emerge in a patchier manner in the early ancestors of the family. These are mostly retained in descendant clades with some losses (Figure 3B). Residues which contact DNA by making specific base contacts clearly emerge at the overall common ancestor and are strongly conserved downstream (Figure 3C).

**Figure 3:**
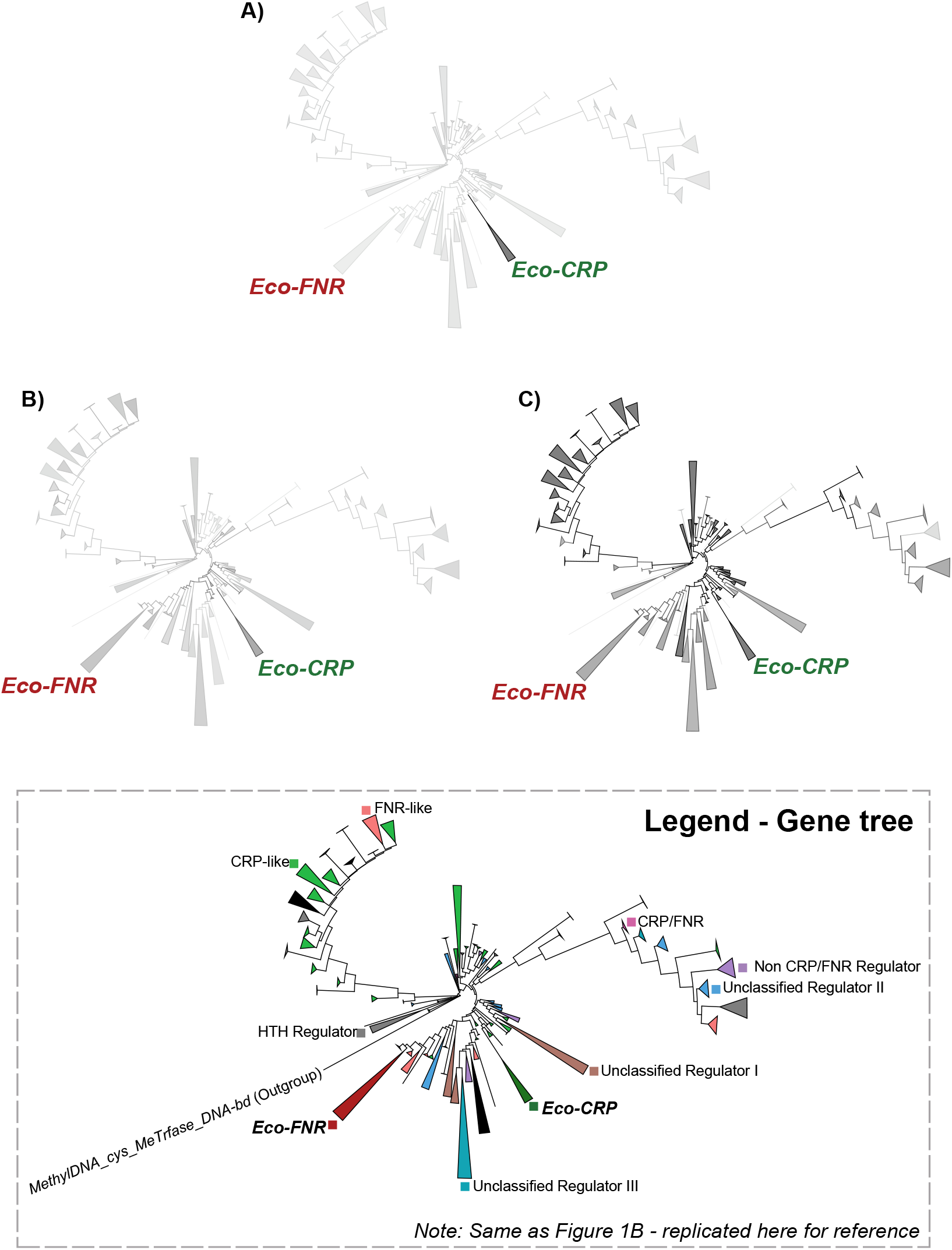
Conservation and emergence in the gene tree. Collapsed outgroup rooted gene trees of the c-f-HTH domain (N=10,224) coloured on a grayscale gradient (low to high) on the basis of PSSM scores calculated for (A) Activating Region 1 (AR1) residues, (B) Residues involved in making phosphate backbone contacts, (C) Residues involved in making specific base contacts with DNA. Size of collapsed nodes corresponds to the number of extant tips. Legend in grey box is Figure 1B replicated for reference to identity of different clades in the gene tree. Collapsed and visualized using iTOL

Thus, in the c-f-HTH domain, DNA binding by base specific contacts likely emerged in the overall family common ancestor. The phosphate backbone contacts, present minimally in the overall family ancestor, gain conservation in a patchy manner across the gene tree. In contrast, the AR1 site, responsible for contacting the αCTD subunit of RNA polymerase and activating transcription, emerged specifically in the common ancestor of the Eco-CRP clade.

### Possible coevolution of the AR1-αCTD interface

The phylogenetically restricted presence of AR1 presents a few different possibilities: (a) the class I activation through the c-f-HTH domain is a property that is almost exclusive to Eco-CRP; (b) other classes of CRP/FNR family proteins use different interfaces to facilitate class I activation. To test these hypotheses, we analyzed the conservation and possible co-evolution of AR1 and the corresponding interface on the αCTD subunit of RNA polymerase (called the ‘287 determinant’).

We calculated PSSM (Position Specific Scoring Matrix – a measure of conservation) scores for the AR1 residues and compared them to their corresponding scores for the 287 determinant on αCTD. High PSSM scores for both AR1 and the 287 determinant is a property of only Eco-CRP-containing genomes. Whereas AR1 scores are poor for all other clades, the 287 determinant scores vary, suggesting the possibility of alternative AR1 sites in these clades (Figure 4A).

**Figure 4:**
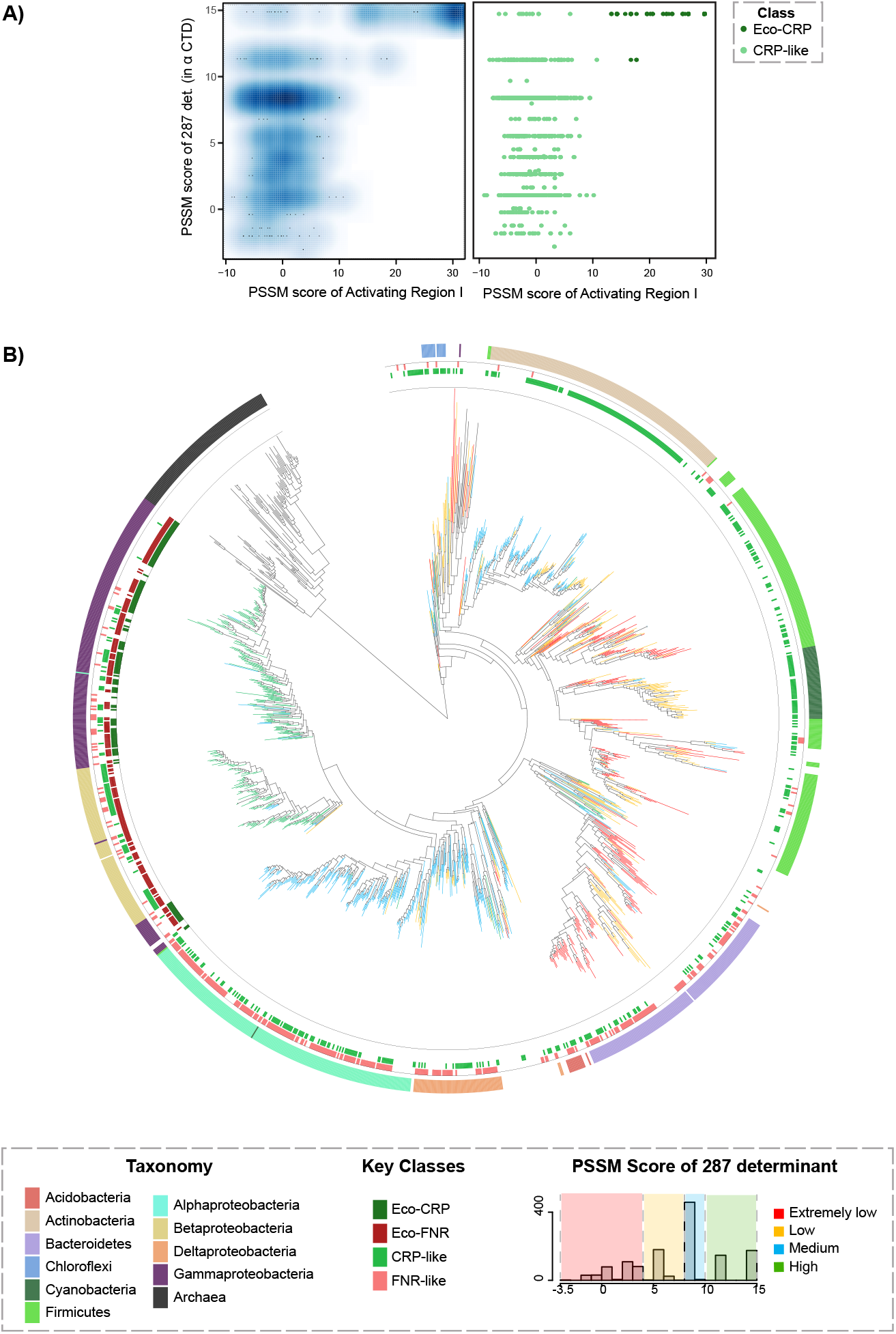
Patterns of conservation of αCTD. (A) Smoothscatter plot and scatterplot of PSSM scores corresponding to 287 determinant of αCTD versus AR1 scores of Eco-CRP and CRP like regulators. The density of the points are shown in the smooth scatter plot while the points are coloured on the basis of class in the scatter plot. (B) Conservation of the 287 determinant mapped onto the 16S rRNA species tree (N=1,454). PSSM scores of the 287 determinant divided into 4 categories as indicated in legend - extremely low, low, medium and high, mapped onto their corresponding genomes on the species tree. Colourstrip annotations show key classes present and the taxonomic annotation of the genome. Note, for simplicity, extremely low and low categories combined into one in the text. Coloured and visualized using iTOL.

**Figure 5:**
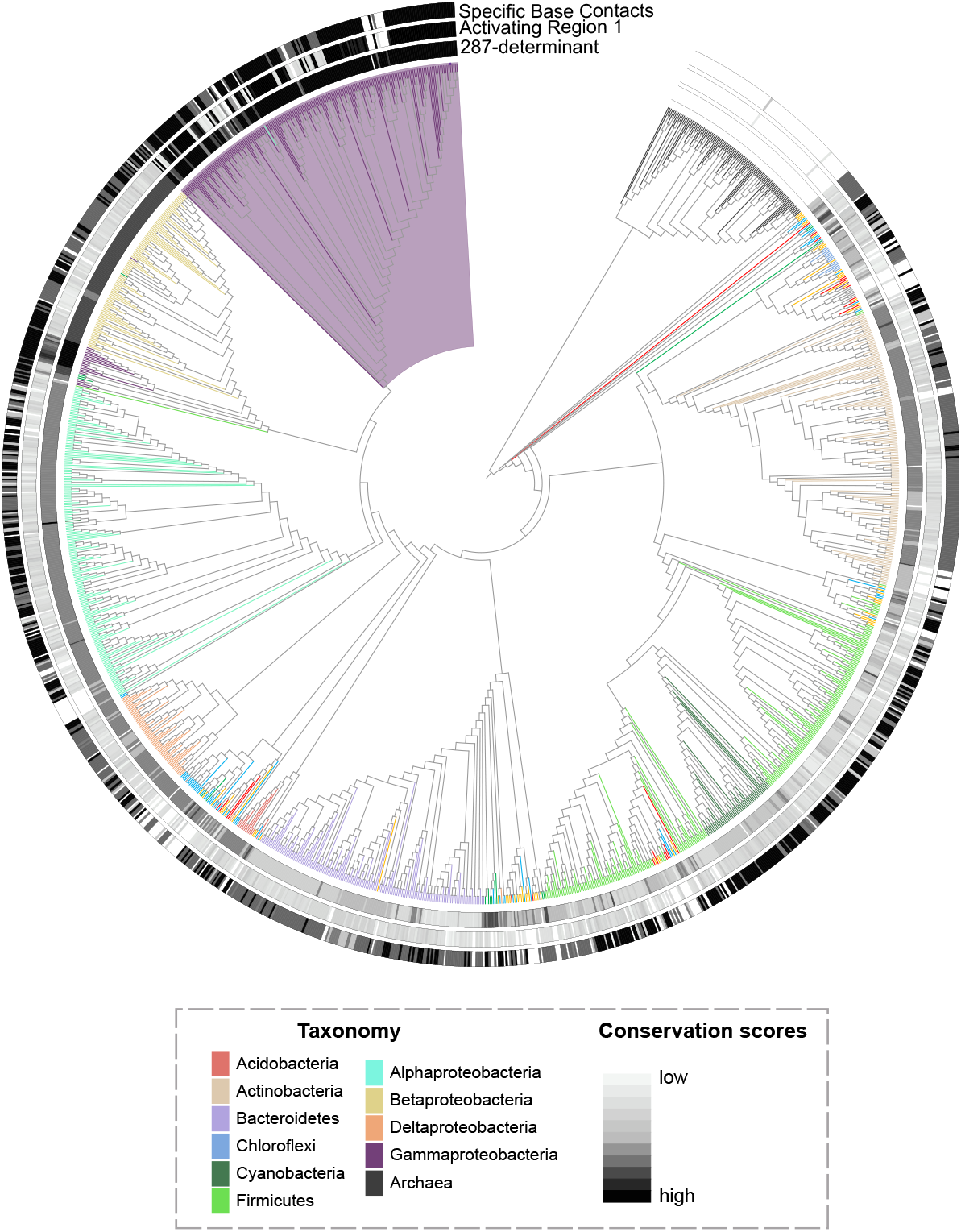
The evolutionary trajectory of functions in the c-f-HTH domain of the CRP/FNR family. Conservation of the key residues identified in the study, marked the 16S rRNA species tree. Colourstrip annotations (innermost to outermost) correspond to the range in PSSM scores observed for the 287 determinant, AR1 and Specific Base contact residues. For genomes coding for multiple CRP/FNR family proteins, AR1 and specific base contact scores for the best-scoring protein are shown. Branches are coloured on the basis of taxonomic annotation, and Gammaproteobacteria, showing the highest scores for 287 determinant of αCTD, along with AR1 and Specific base contact scores is highlighted. Coloured and visualized using iTOL.

In *E. coli* FNR for example, it has been speculated, for the c-f-HTH domain, that an alternative AR1 site, partially overlapping with *E. coli* CRP AR1 may form an interface with αCTD (Lee et al., 2000; S. M. Williams et al., 1997). In FNR, the ligand binding domain also presents a large interface for class I activation, something that has not been described for CRP. We used in-silico modeling to compare the contribution of c-f-HTH domain alone to the interface of CRP and FNR with αCTD (see Supplementary Methods). For CRP, we used the available PDB structure of the *E. coli* CRP-DNA-αCTD complex, from which we extracted only the c-f-HTH domain and αCTD. Due to the unavailability of PDB structures of *E. coli* FNR, we used AlphaFold (Jumper et al., 2021; Varadi et al., 2024) to generate the structure of its c-f-HTH domain. To test the interface, we performed a docking simulation using HADDOCK 2.4 (Honorato et al., 2024) on both the control CRP-αCTD and the test FNR-αCTD structures, providing the proposed AR1 and 287 determinant residues, as well as the residues identified by (S. M. Williams et al., 1997), as the active sites. Of all the complexes produced by docking of *E. coli* CRP and αCTD, nearly three-fourths (N = 106 / 146) produced the expected interface. In contrast, only a third of the docked structures (N = 24 / 80) for *E. coli* FNR-αCTD produced any interface, and this was not even the predicted interface (Supplementary Figure 4). This suggests that, unlike in *E. coli* CRP, the c-f-HTH domain in FNR alone probably does not form a stable interface with αCTD using AR1.

To assess the possible presence of alternative AR1 sites in different CRP clades, we hypothesized that such novel and functional AR1 patches, if present, might be locally conserved in some clades. We first assembled class-specific c-f-HTH domain sequence alignments of all 10 classes of proteins. These showed that the patch contributing to DNA binding is conserved in all classes, but other than Eco-CRP, only Eco-FNR shows a conserved patch that could contribute to a putative FNR-AR1 (Supplementary Figure 5A, B). Since CRP-like is a large and diverse class, we looked at clade-specific alignments of two distinct CRP-like clades: a CRP-like clade closer to the ancestor in the gene tree (termed “Anc-CRP-like”) and another CRP-like clade which is a sister clade to Eco-CRP (termed “Sister-CRP-like”). Clade-specific alignments of the Anc-CRP-like and Sister-CRP-like also show some possible conserved patches outside of the DNA binding patch (Supplementary Figure 5C,D), which could serve as putative AR1 residues; however, the degree of conservation, as revealed by within-clade PSSM scores, is relatively low, pointing to possible lack of AR1-like functionality (Supplementary Figure 6).

We then turned to looking at patterns of conservation of the second component of the interface: the 287 determinant of αCTD, using a PSSM constructed from the *E. coli* protein. We built an unrooted α CTD gene tree (see Supplementary Methods) for 1,454 genus level curated taxa (Supplementary Figure 7). We however faced issues in choosing a suitable outgroup for rooting this tree, as it is ubiquitously present across all Bacteria, with no other members present in the homologous superfamily of the domain. Further, the unrooted αCTD tree contained some extremely long branches. Given these issues, under the reasonable expectation that the gene tree of a core component of the central dogma would largely track the species tree, we chose to examine the evolutionary trajectory of αCTD, in particular that of the 287 determinant, on the species tree (Figure 4B). We found that the 287 determinant is poorly conserved i n a lmost a l l *Firmicutes* and *Cyanobacteria*. A*ctinobacteria*, along with most classes of *Proteobacteria* (except *Gammaproteobacteria*) show medium conservation; however, the two distinct clades of bacteria show different sets of conserved residues. This indicates that partial evolution of the 287 determinant started independently in two distant clades: *Actinobacteria* and *Proteobacteria* at their respective common ancestors. No further progress appears to have happened in *Actinobacteria*. However, in the *Proteobacterial* lineage, additional residues appeared at the common ancestor of *Beta*- and *Gammaproteobacteria*. All residues comprising the 287 determinant emerged at the common ancestor of *Gammaproteobacteria*, to which *E. coli* belongs (Supplementary Figure 8). Hence, the evolution of the 287 determinant as seen in *E. coli*, started from the *Proteobacterial* ancestor and became fully formed in *Gammaproteobacteria*, indicating a stepwise evolutionary trajectory for the conserved surface in αCTD.

Thus, we find that the interface between c-f-HTH and αCTD shows a high possibility of coevolution, with a highly conserved 287 determinant found only in conjunction with a conserved AR1 only in the Eco-CRP clade present in *Gammaproteobacteria*. While other interfaces could exist, we show that these possibilities are not apparent in conservation patterns in the c-f-HTH or in αCTD.

## Discussion

The CRP/FNR family is a conserved family of TFs with a well-characterized ligand binding domain and a helix-turn-helix-containing DNA binding domain. A paper from over 20 years ago (Körner et al., 2003) provided a comprehensive coverage of the different classifications of CRP/FNR TFs found in prokaryotes, but was limited by the number of available genomes at that time. More recent studies (Matsui et al., 2013) have attempted to look at the evolutionary history of the CRP/FNR family. Recognizing the difficulties in constructing an outgroup-based phylogeny of the CRP/FNR family, Matsui and colleagues used a method called spectral clustering to classify members of the family to indirectly infer ancestral states. They predicted that the ancestral protein could be more similar to FNR, and that CRP proteins emerged later. Our rooted phylogenetic analysis suggests an alternative hypothesis, where the early ancestral DNA binding domains of the family are more similar to members of a large and diverse class of “CRP-like” proteins, and distinct from close orthologs of *E. coli* CRP (Eco-CRP) and FNR (Eco-FNR). This discrepancy in the nature of the early ancestor could possibly be attributed, besides methodological differences, to the fact that Matsui et al. had analysed full protein sequences, whereas we investigated only the c-f-HTH domain. The ligand binding domain could have taken an independent evolutionary trajectory, ultimately influencing the whole protein evolution.

We also show that while residues conferring DNA binding specificity are present from the overall common ancestor, binding to RNA polymerase via the Activating Region I (AR1) mechanism is specific and restricted to Eco-CRP. AR1 is essential for Class I type of transcription activation and has been extensively studied in *E. coli* CRP (Ebright, 1993). A comprehensive review of *E. coli* CRP, examining the structure, allostery, transcriptional activation and repression of CRP in extensive detail, compared its DNA binding domain with 12 of the most well characterized members of the family available at the time (Kolb et al., 1993). This study indicated that the surface exposed loop of AR1 is absent in other regulators from the family and appears to be found uniquely in *E. coli* CRP. While their study largely focuses on CRP and FNR from *Escherichia coli*, other members reviewed belong to *Gammaproteobacteria* and *Alphaproteobacteria*, with 2 members belonging to *Cyanobacteria*, and upon comparison, most of these proteins fall in the FNR-like category of our dataset and thus do not represent the diversity of the current dataset. Using our expanded dataset, we show that the AR1 loop is unique to the Eco-CRP clade, and has likely evolved in its own common ancestor.

Most of the literature on this family is concerned with *Escherichia coli* CRP. However, a recent review (Krol et al., 2023) states that DNA binding properties may be conserved across various experimentally-investigated members of the family f rom *Proteobacteria* and *Cyanobacteria*. A ‘non-exhaustive’ phylogenetic analysis (Soberón-Chávez et al., 2017) provides insights into the members of this family across multiple taxa, mostly belonging to the large phyla of *Proteobacteria* and *Actinobacteria*, concluding that there is ‘some’ conservation of DNA binding specificity. Thus, consistent with our analysis, the core DNA binding motifs remain conserved across taxa. However, there is significant variation in mechanisms of transcriptional activation or repression. To our knowledge, AR1 has not been experimentally defined in outside Eco-CRP. For example, there exists a hypothesis that CRP and FNR in *E. coli* may possess entirely different interface residues with αCTD to facilitate Class I activation (Lee et al., 2000). However, as noted in the above study, this Class I FNR-dependent transcription activation complex could not be verified in vitro. Our sequence analysis also does not suggest the presence of a well-conserved AR1 in other classes of the family.

Several members of this family appear to use Class II modes of activation. This has been studied in detail in FNR, particularly in examples in which FNR acts in conjunction with other redox sensing regulators such as NarL and NarP to coregulate gene expression (Mazoch & Kučera, 2002; Myers et al., 2013). Note here that AR1 also contributes to Class II activation in *E. coli* CRP (Rhodius et al., 1997; West et al., 1993). A structural analysis of a Class II activation complex formed by a CRP/FNR family regulator in *Thermus thermophilus* (Feng et al., 2016) shows that the AR1 residues predicted using *E. coli* CRP do not form an interface with αCTD but instead, a larger surface termed “AR4” contributed by both the domains contacts αCTD, at residues other than the 287 determinant (at the extreme C terminal end of αCTD). All the prior studies combined with the results from our own analysis indicates that the emergence of AR1 in the c-f-HTH domain is likely an independent gain in the *Gammaproteobacterial* common ancestor. This raises the question of whether class I activation, involving only one interface of CRP with αCTD, is a relatively weaker mode of activation compared to class II activation, which requires multiple contacts of the TF with the body of the RNA polymerase. If so, given that class I activation can be flexible in terms of the relative positioning of the TF binding site and the promoter, has it evolved to facilitate integration of potentially sub-optimal promoters into the global regulatory network of *E. coli* CRP?

Although our analysis in this study is restricted to the c-f-HTH domain, we wish to highlight the roles played by the ligand binding domain in RNA polymerase interactions (Berman et al., 2005). The predicted AR1 residues for FNR in *E. coli* are spread out into 3 surface-exposed loops, one of which, present in the ligand binding domain (116-121), is important for Class I activation by possible contacts to the extreme C-terminal end of αCTD (R. Williams et al., 1991). However, this loop in the ligand binding domain of CRP does not play a role in CRP-mediated Class I activation. Further analysis of the ligand binding domain independently could elucidate the role of this loop in the other classes of the CRP/FNR family of TFs.

The αCTD of RNA Polymerase evidently plays a clear role in transcriptional activation facilitated by CRP, and our analysis indicates that the set of residues which interact with CRP in *E. coli* might have co-evolved with AR1. However, while AR1 appears to have abruptly evolved in *Gammaproteobacteria*, the 287 determinant started to form in the *Proteobacterial* common ancestor, becoming better established in the *Gamma*- and *Betaproteobacterial* ancestor before being fully formed in its extant state in *Gammaproteobacteria*. Whether AR1 drove the final formation of the Gammaproteobacterial 287 determinant or vice-versa could not be determined. To our knowledge, the hypothesis that some activators may contact αCTD at its extreme C-terminal residues, remains to be validated (Lamberg & Kiley, 2000; S. M. Williams et al., 1997), due to a larger focus on activator-DNA contacts compared to activator-αCTD contacts.

In summary (Graphical Abstract), we show that in the CRP/FNR family of TFs, the interface between CRP and αCTD in the Class I activation is unique and phylogenetically constrained only to Eco-CRP, while DNA binding is ubiquitous across the c-f-HTH domains in the CRP/FNR family.

## Data Availability

All necessary data is available at https://doi.org/10.6084/m9.figshare.29476913.v1. Supplementary materials available to download.

## Acknowledgements

We thank Sunil Laxman, Inder Raj Singh, Ashitha B Arun, Ganesh Muthu, Pragya Tiwary and Akshara Dubey for discussions on the project.

