## Supplementary Figures and Supplementary Methods for "The evolution of function in the DNA binding domain of the CRP/FNR family"

### Supplementary Materials

#### Supplementary Figures

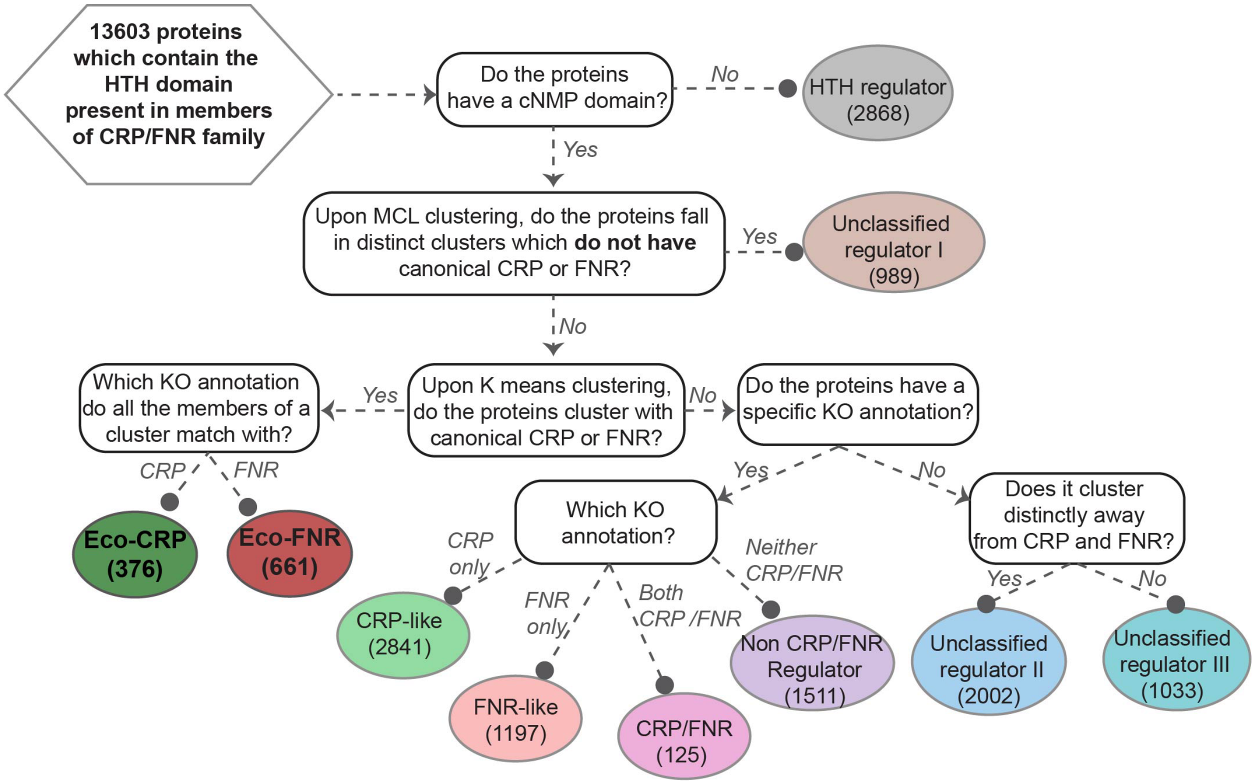


**Supplementary Figure 1: Orthologous classification of CRP/FNR family of regulators.**

Flowchart detailing the different steps and checkpoints used for classification of orthologous groups in CRP/FNR family. A combination of clustering algorithms based on similarity scores (RBH, mcl, Kmeans) along with KO annotations used to arrive at the final number of classes (10).

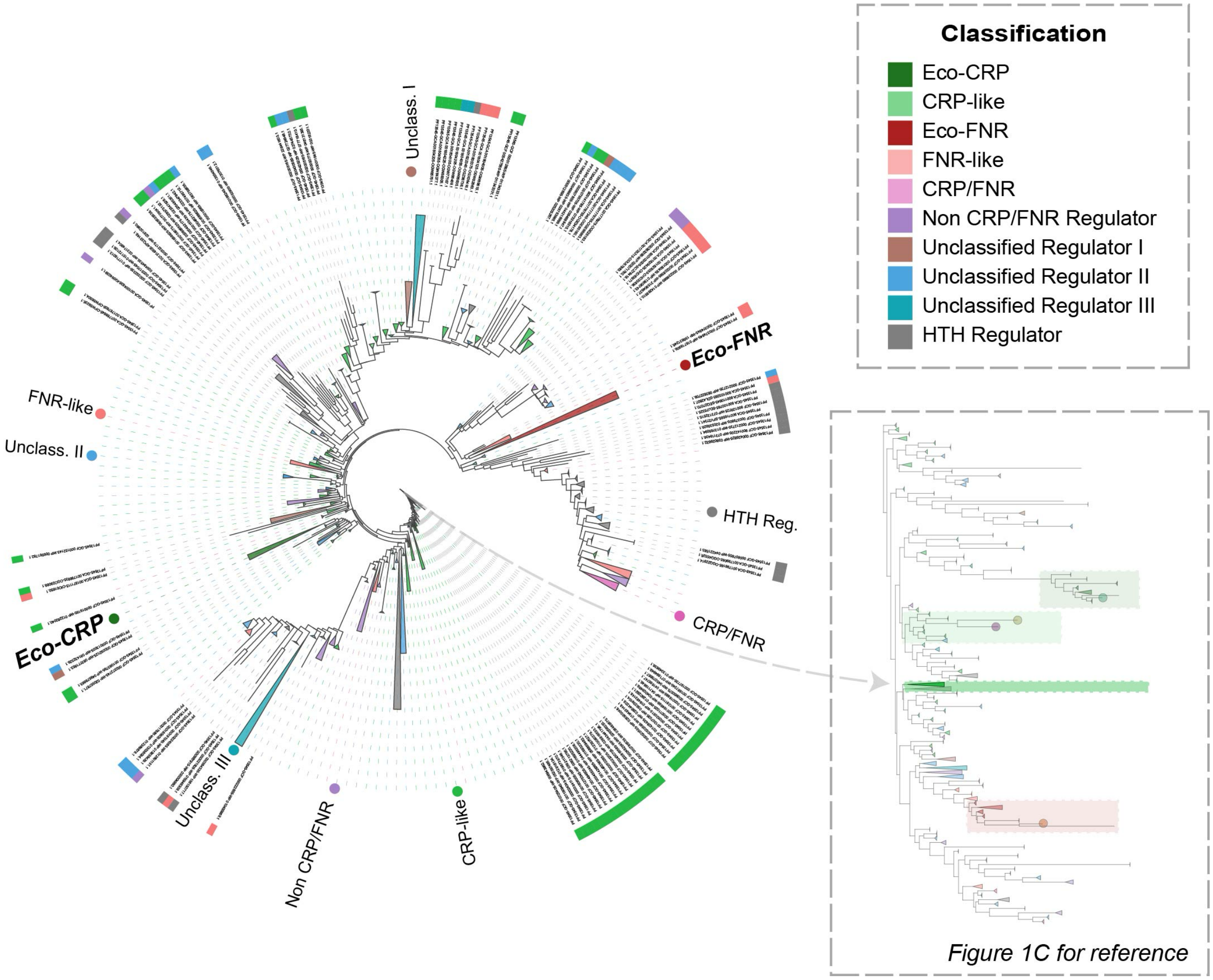


#### Supplementary Figure 2: Optroot rooted gene tree of c-f-HTH domain

Partially collapsed rooted gene tree (N=10,223) of c-f-HTH domains. The optimal rooting (based on a CRP like protein) was identified from the domain sequences present in the alignment on the basis of phylogenetic reconciliation. Size of collapsed nodes corresponds to number of extant tips, and coloured based on protein classification established in Supplementary Figure 1. Collapsed and visualized using iTOL. Inset of Figure 1C highlights the clade which contains the CRP-like sequence identified from the optimal rooted tree.

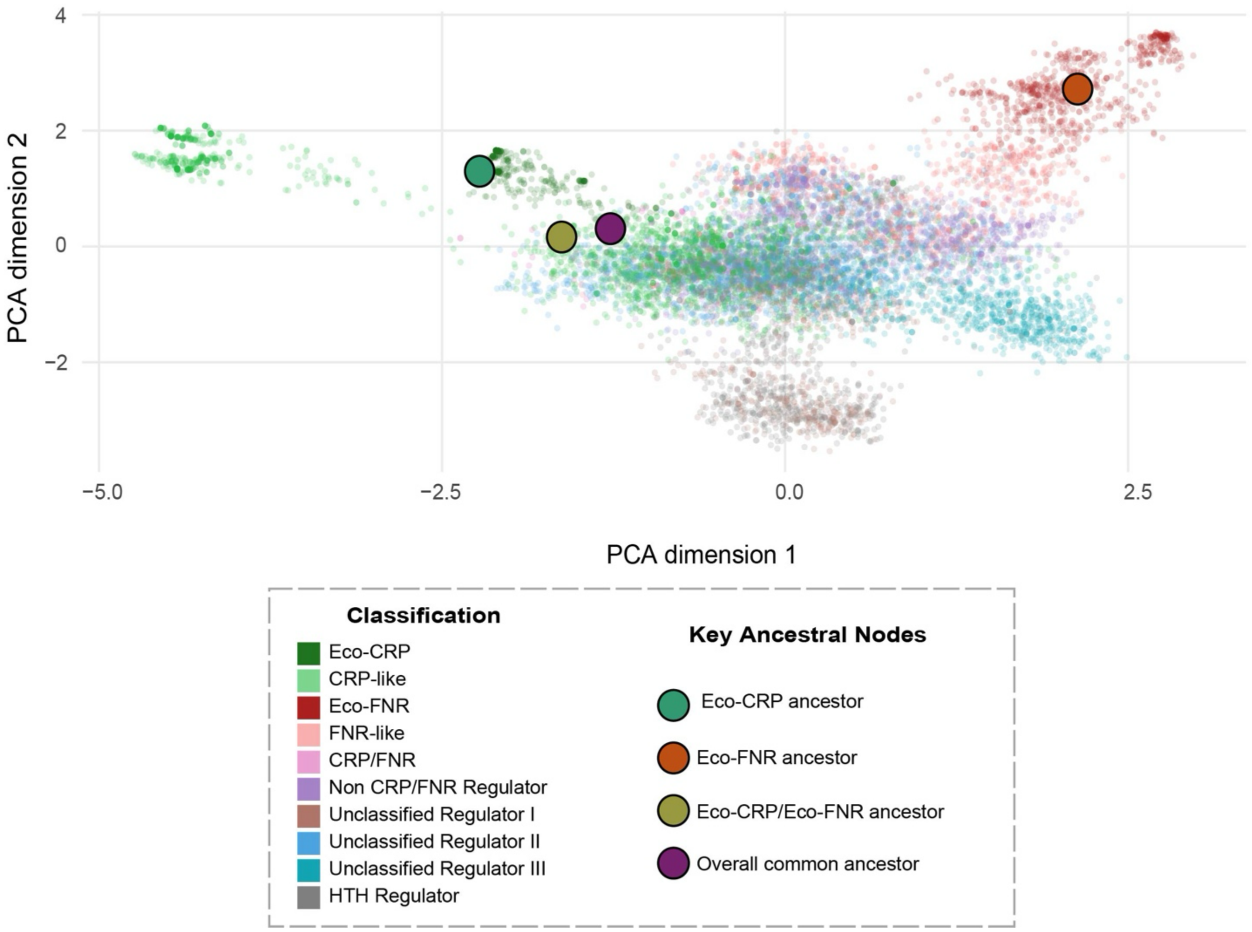


**Supplementary Figure 3: Principal Component Analysis (PCA) of c-f-HTH domains with key ancestral nodes**

PCA scatterplot constructed from the c-f-HTH domain alignment with key ancestral nodes, coloured on the basis of the protein classification. Key nodes highlighted and included the overall common ancestor of the family (purple), the common ancestor of Eco-CRP and Eco-FNR (yellow) and corresponding common ancestors of Eco-CRP (green) and Eco-FNR (red).

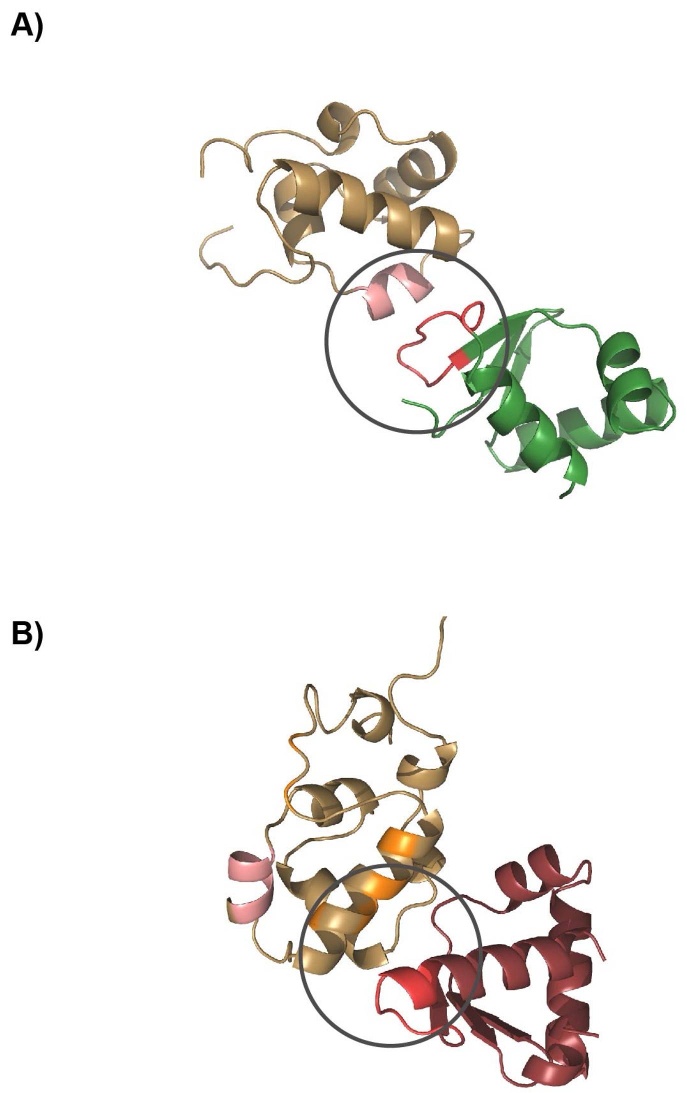


**Supplementary Figure 4: Docked complexes of CRP and FNR with αCTD in *E. coli***

Docking results for the Class I activation complex in *Escherichia coli* provide a number of water refined structures, sorted into clusters, which are scored based on their average RMSD and energetic properties. An average structure is generated for each cluster. The average structure of the highest scoring clusters for the control and test runs are shown in (A) and (B), with αCTD coloured sand yellow. (A) the c-f-HTH domain of CRP (control-coloured green) with αCTD (B) The c-f-HTH domain of FNR (test - coloured red). The AR1 region as found in CRP is coloured red, and the corresponding 287 determinant in αCTD is coloured pink, with the extreme residues of αCTD coloured orange.

Interface predicted by docking analysis highlighted with a grey circle in both complexes. Coloured and annotated using PyMOL.

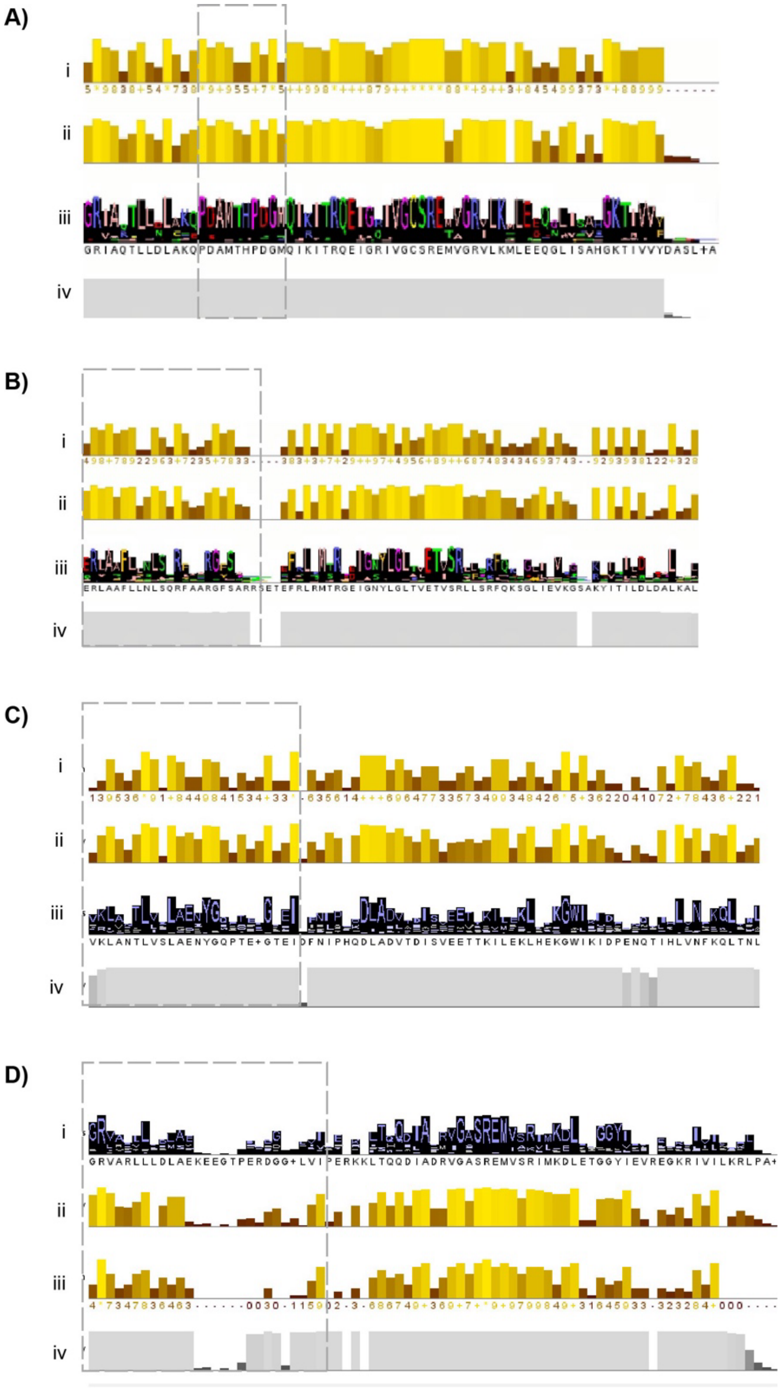


**Supplementary Figure 5: Class specific and clade specific alignments of c-f-HTH domains**

Class specific alignment scorings of the (A) Eco-CRP class, (B) Eco-FNR class, (C) Anc-CRP-like clade, (D) Sister-CRP-like clades. Putative AR1 patches highlighted in the grey dashed box. (i) indicates the conservation score (0-11) for each residue in the alignment, (ii) corresponds to the quality of the conserved site, (iii) indicates the consensus (major amino acid frequency) for a residue site, (iv) gives an extent of coverage (occupancy) at each site. These scorings are obtained from Jalview.

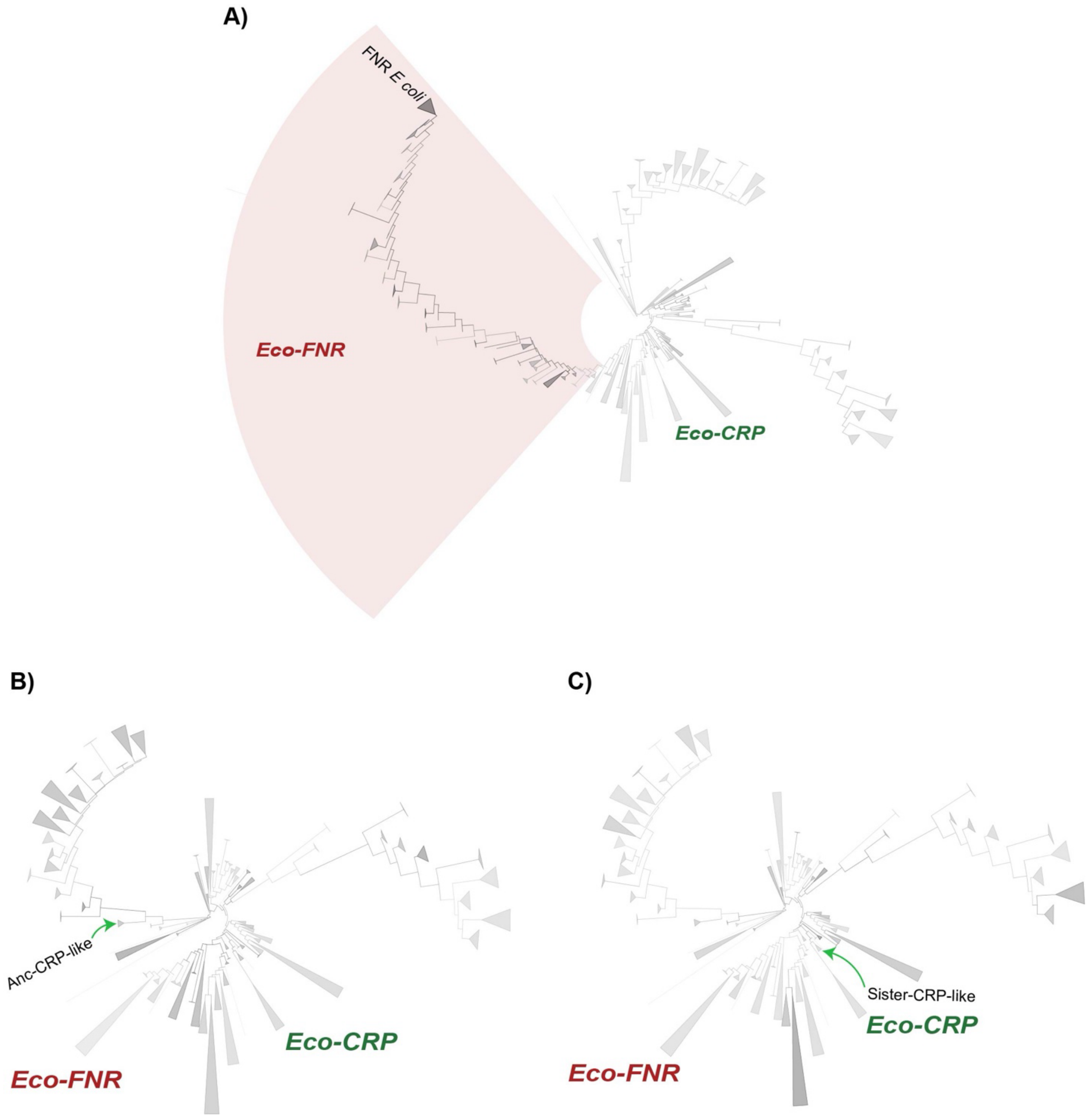


#### Supplementary Figure 6: PSSM scores of putative AR1 residues

Collapsed outgroup rooted gene tree (N=10,224) coloured on a grayscale gradient (low to high) on the basis of PSSM scores calculated for (A) putative FNR AR1 residues, (B) putative Anc-CRP-like AR1 residues and (C) putative Sister-CRP-like AR1 residues. The FNR clade was expanded from the earlier view in Figure 1B for (A), and subcollapsed accordingly, with all clades belonging to the FNR clade shaded in red. The corresponding CRP-like clades have been marked with an arrow in (B) and (C). Remaining collapsed nodes consistent with earlier gene trees, collapsed and visualized using iTOL.

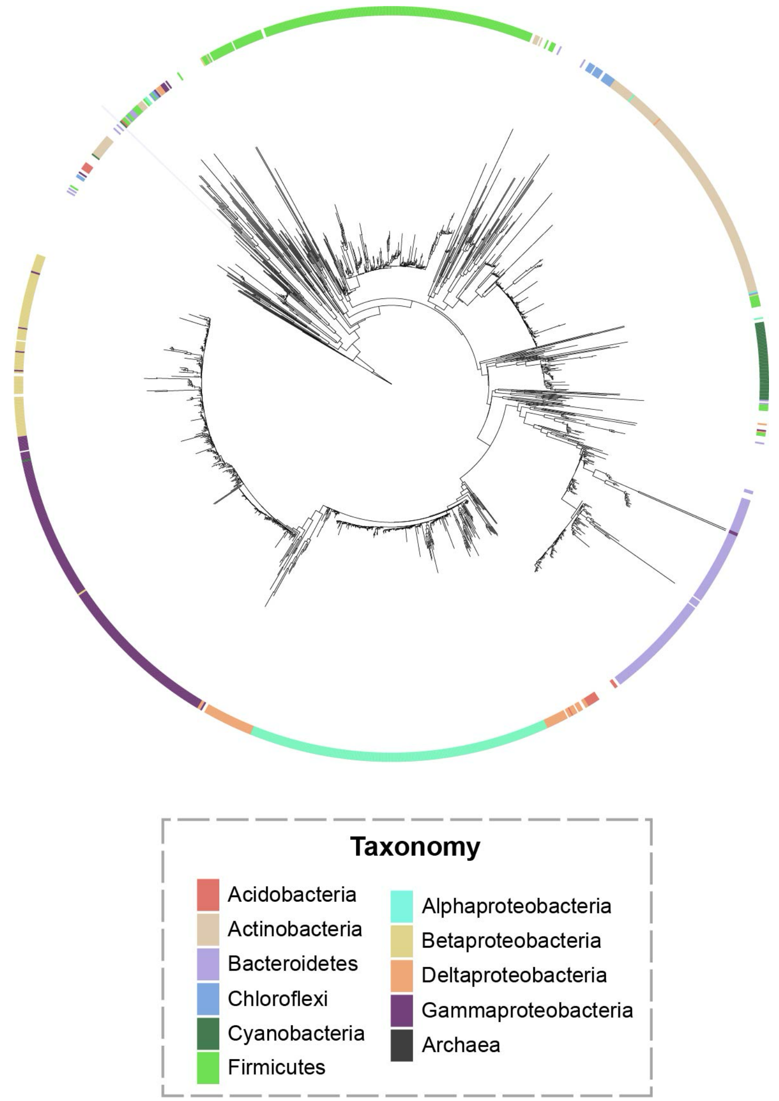


#### Supplementary Figure 7: The αCTD tree

Unrooted gene tree (N=1,442) of αCTD gene sequences assembled from 1,328 taxa (~120 Archaea in dataset of 1,454 taxa do not contain αCTD sequences) . Colourstrip annotation refers to the corresponding taxonomic annotation of the αCTD sequence. Visualized using iTOL.

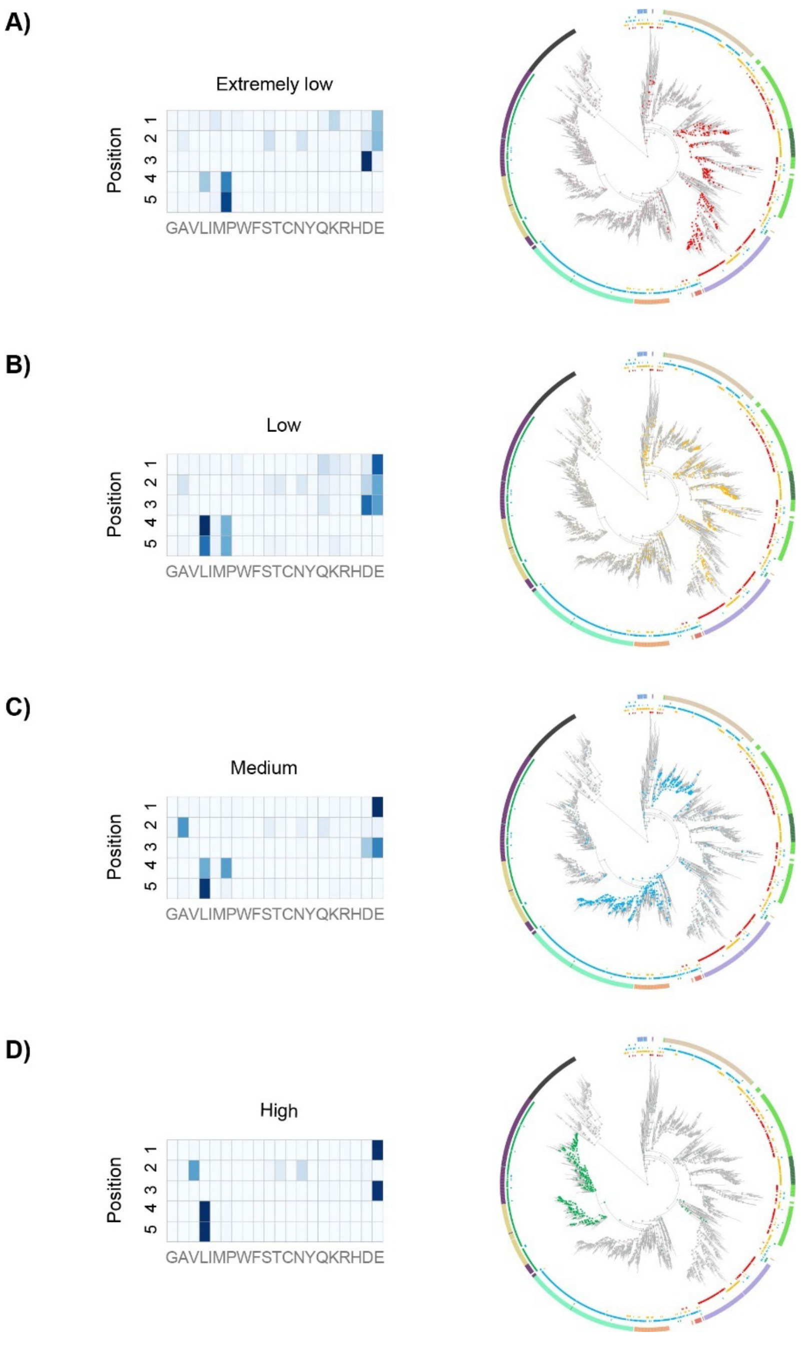


**Supplementary Figure 8: Ancestral state reconstruction of PSSM scores on the species tree**

Ancestral state reconstruction of the levels of conservation identified for the 287 determinant, done on the 16S rRNA species tree (N=1,454), along with the composition of residues, for (A) Extremely low PSSM values (<3.2) (B) Low PSSM values (between 3.2 and 8.4), (C) Medium PSSM values (between 8.4 and 10) and (D) High PSSM values (>10). For reference, the score for *E. coli* is 14.6. The heatmap is coloured from light (low) to dark(high) for conservation of a particular amino acid at the respective position. Trees are visualized and annotated using iTOL.

### Supplementary Methods

#### Protein curation

To obtain orthologous members of the CRP/FNR family of proteins, hmmsearch using hmmer3 was performed for the ligand binding cNMP domain of canonical CRP (Pfam: PF00027), and the DNA binding HTH domain of canonical CRP (Pfam:PF13545) (referred to as the c-f-HTH domain), with a trusted cutoff set to remove weakly scoring hits. The obtained hits were used for a hmmscan against the entire Pfam profile database to obtain complete domain architecture. Since domain annotations often overlap with multiple hits for the same residue coordinates, the two domains of interest (if present) were taken as primary annotation, and if unavailable, the domain annotation corresponding to the longest stretch of residues was taken to ensure maximum coverage of the protein. The final list of proteins belonging to the CRP/FNR family consisted of proteins with the two non-overlapping domains (N=10,736). For further analysis involving only the c-f-HTH domain, the proteins which had the c-f-HTH domain without the ligand binding domain were included, to give a total of 13,603 proteins.

#### Orthologous classification

First, proteins with only the c-f-HTH domain and without the ligand binding domain were classified as HTH regulators. Subsequently, an all against all BLAST (blastp) (Camacho et al., 2009) was performed against whole protein sequences containing both domains of interest, with an e-value cutoff of $10^{-3}$. The hits obtained above the e-value cutoff were taken and a reciprocal best hit was performed based on the bitscores. The RBH pairs were then clustered using a Markov Cluster Algorithm or "mcl" (van Dongen & Abreu-Goodger, 2012), on the basis of the bitscores. The stringency of the clustering is controlled by the inflation parameter I, found to be 1 for this dataset by the elbow method.

To resolve ambiguity with the clusters obtained from mcl, additional methods of classifying the proteins had to be employed. A Principal component analysis (see below) of the protein sequence alignment was performed, and the obtained principal components were then clustered based on Kmeans clustering. To supplement the clustering results with functional annotations, the proteins were run against eggNOG-mapper to retrieve KO ids of CRP and FNR, along with (if present) KO ids of other members of the CRP/FNR family. A combination of the functional annotations along with the clustering with respect to CRP and FNR of Escherichia coli were used for the final classification into 9 orthologous groups of CRP/FNR, with an additional HTH regulator group.

#### Pruning and curation of domain sequences

First, a length cutoff was set for each sequence in the alignment based on the distribution of lengths for the domains, removing outliers that were too long or too short. Following this, the subset of sequences was realigned and systematically pruned on the basis of gaps (sequences with >5 residues contributing to >95% gaps removed) and a bitscore cutoff (sequences with bitscore <40 removed from alignment).

#### PCA calculation from an alignment

An initial empty matrix is made in which the number of rows corresponds to the total number of sequences in the alignment, and the number of columns corresponds to the probability of a residue present in a particular column position of the alignment, with elements corresponding to 0/1. Given a protein alignment (20 + a gap character) of length m with n sequences, the sequence matrix $S_{ij}$ consists of a n x (21m) logical matrix, where if for sequence 1, residue position 1 has amino acid A, $S_{11} = 1$ and $S_{1j} = 0$ for $1<j<22$.The same is computed for all residues across all sequences, and all the possible values for a particular column position are assembled together. The PCA is run using sklearn, and the two dimensions contributing to the most variance are plotted.

#### Structural analysis of protein complexes

An initial approach was to model structures based on AlphaFold2 using the hetero-oligomer option and with a custom template of the active conformation of a reference complex (Modified CRP-αCTD complex (PDB: 1LB2) with only the two domains). However, these gave inconsistent results with low-confidence models, attributed to low confidence scores and the inherent noisy nature of AlphaFold predictions for complexes. Swiss-Model (homology based) proved to be a better alternative. The modelled structures were used to identify and validate the residues involved in interactions. Docking of a protein-protein complex was conducted using HADDOCK 2.4. The active sites were specified to be the corresponding key residues from literature and predictions, run with all the default parameters for docking a protein-protein complex. Docking results provided the number of clusters identified for each of the runs: for CRP – 146 structures classified into 3 clusters, representing 73% of the water refined models, while for FNR 80 structures classified into 13 clusters, representing 40% of water refined model. The structure from the highest scoring cluster from each run was taken and visualised using PyMOL.

#### Construction of αCTD tree

The same steps adopted for the c-f-HTH domain gene tree construction was also employed for the αCTD gene tree, with the number of genomes subset to the 1,454 genomes used in the 16S rRNA species tree. The αCTD domain was identified on the basis of the corresponding Pfam annotation (PF03118). The tree was constructed without a specified outgroup or asr, due to issues in identifying suitable outgroups for the family, run using IQ-TREE 2 with a "KOSI07+R10" model for 1000 ultrafast bootstrap replicates. The resulting tree (N=1,442) had some long branches which could not be resolved completely along with instances of multiple copies of the domain in the dataset and hence, the 16S rRNA species tree was used for further estimations.

Ancestral state reconstruction on the species tree follows from previous work and ancestral states are taken as discrete characters corresponding to the levels of conservation of the 287 determinant, performed using the *ape* (Paradis & Schliep, 2019) package from R.
